## Supplemental Figures for "The apicoplast link to fever-survival and artemisinin-resistance in the malaria parasite"

**Supplemental Figure S1**

**Schematic overview of pooled phenotypic screening pipeline**

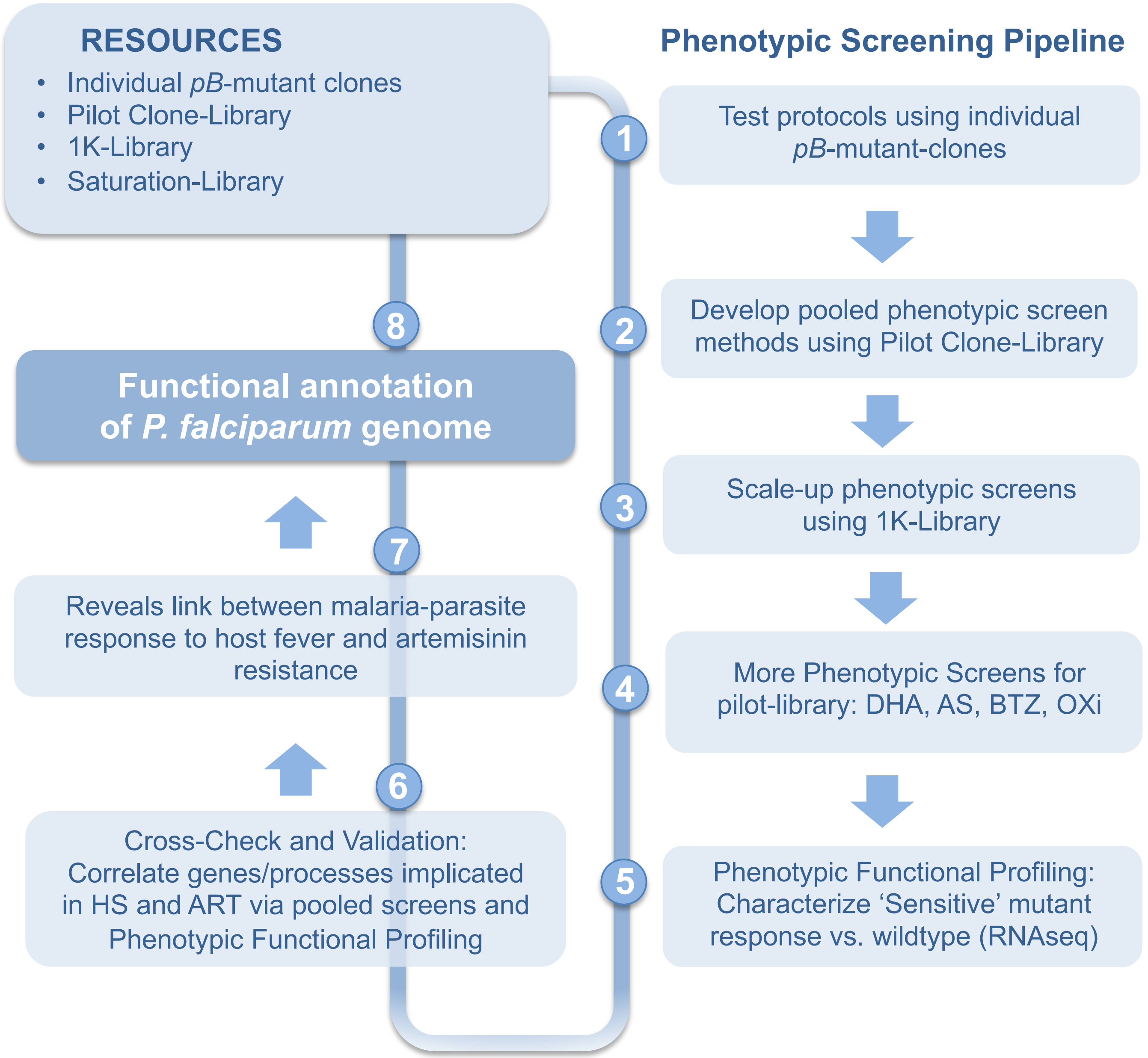

**Supplemental Figure S2**

**Extended phenotypic screening data against the Pilot-library and summary**

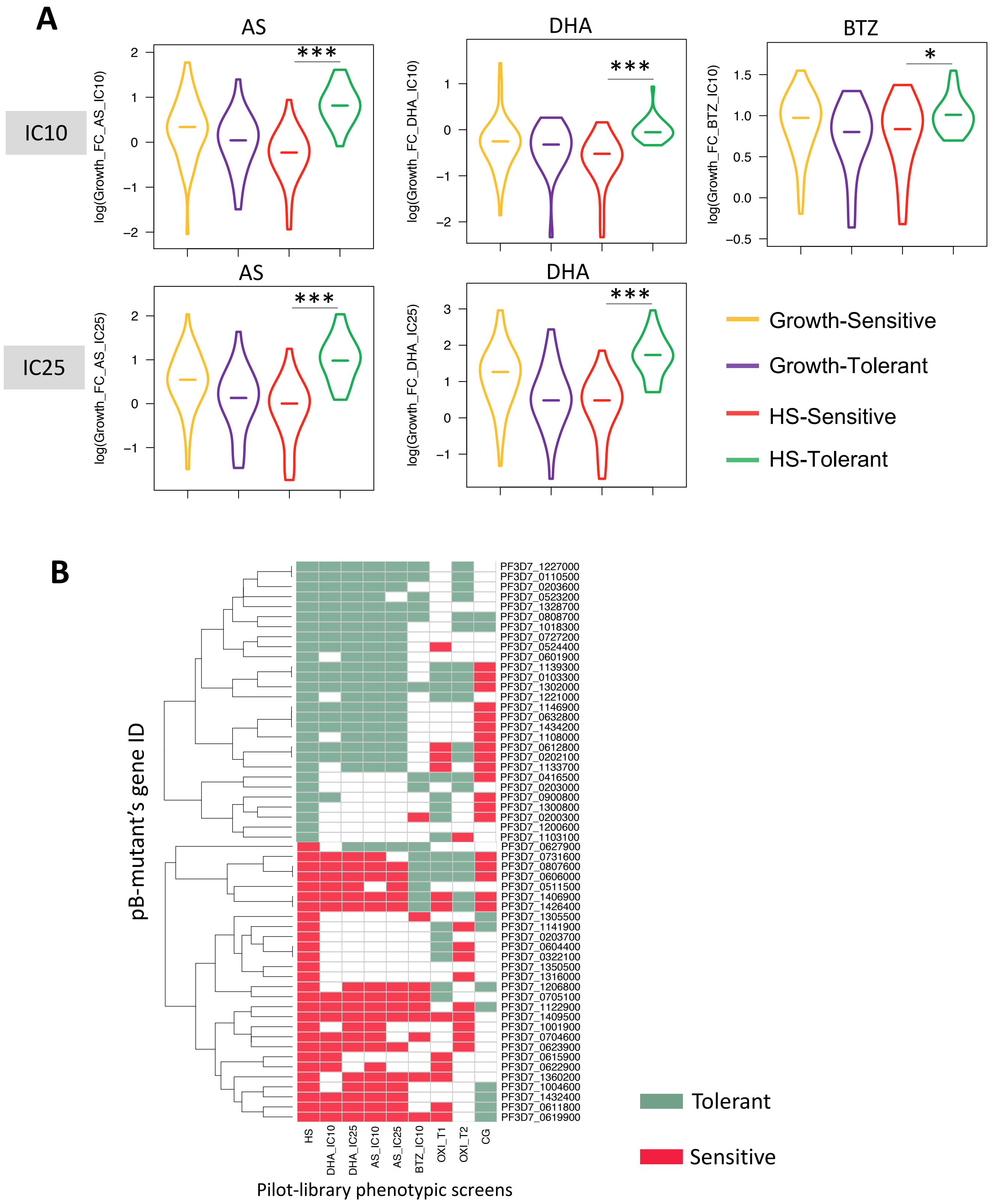

**Supplemental Figure S3.**

**Mutants in (A) members of the DV proteome, (B) targets of ART alkylation, and (C) putative K13-interacting partners tend to be sensitive to HS**

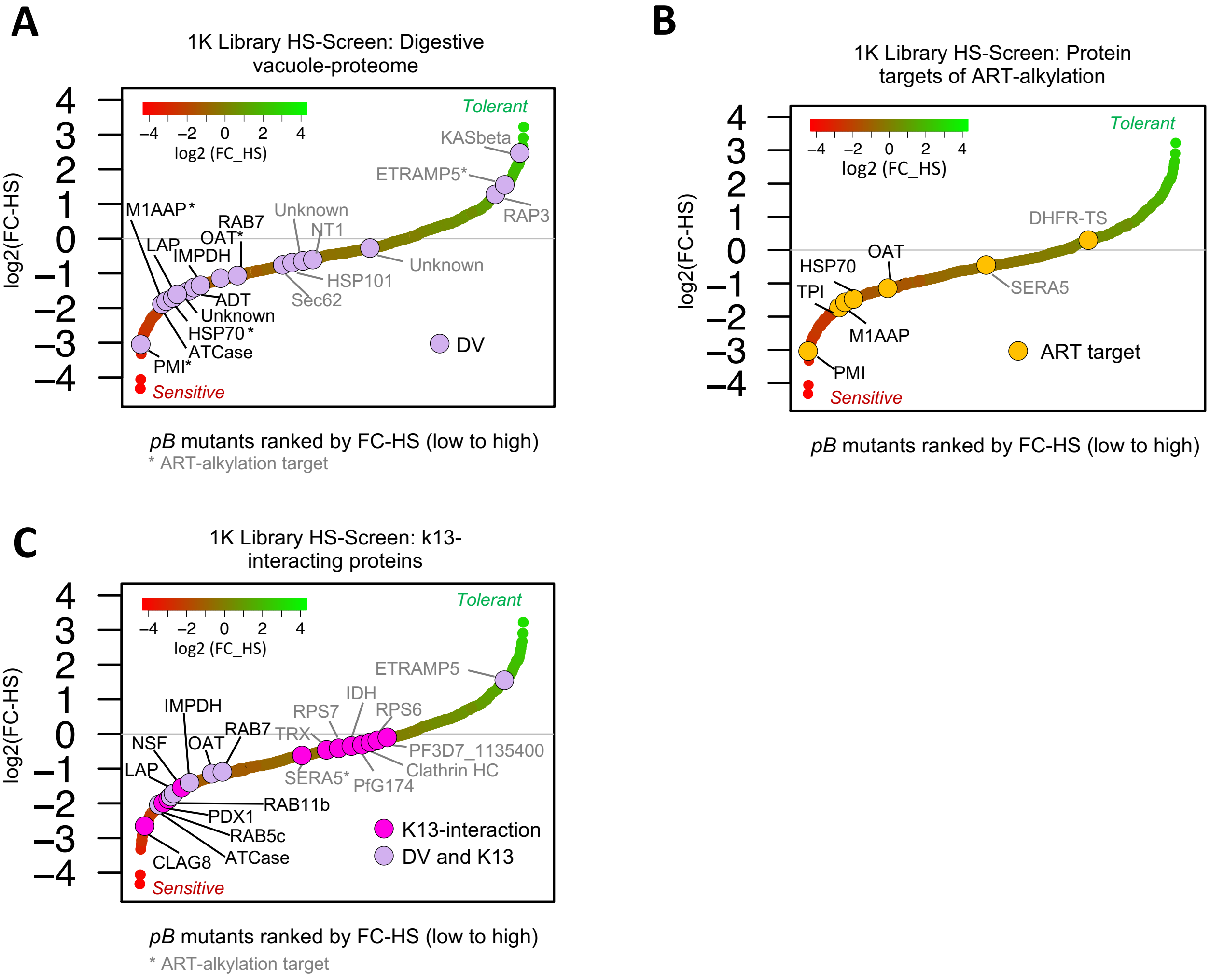

### Supplemental Figure S4.

**A.**

**Core proteasome-components are slightly but universally upregulated in response to HS**

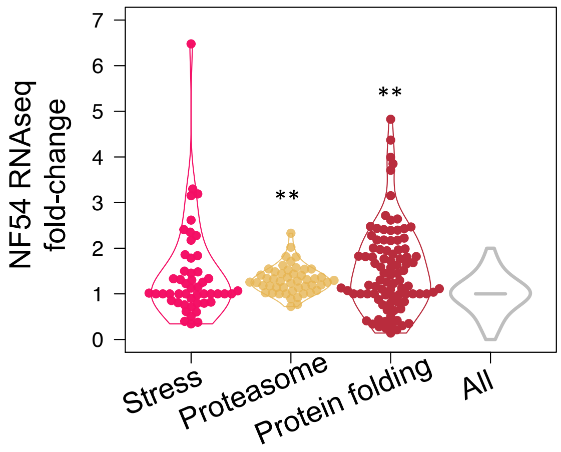


**B.**

**Activation of pathways underlying DHA-mediated killing and febrile-temperature survival is directly inverse**

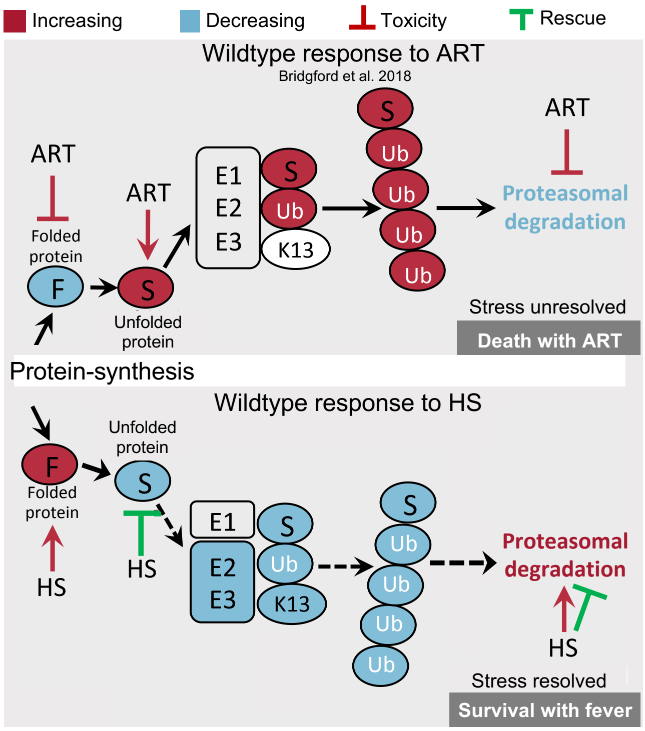

**Supplemental Figure S5.**

**QIseq correlations between biological replicates of Pilot-Library Screens**

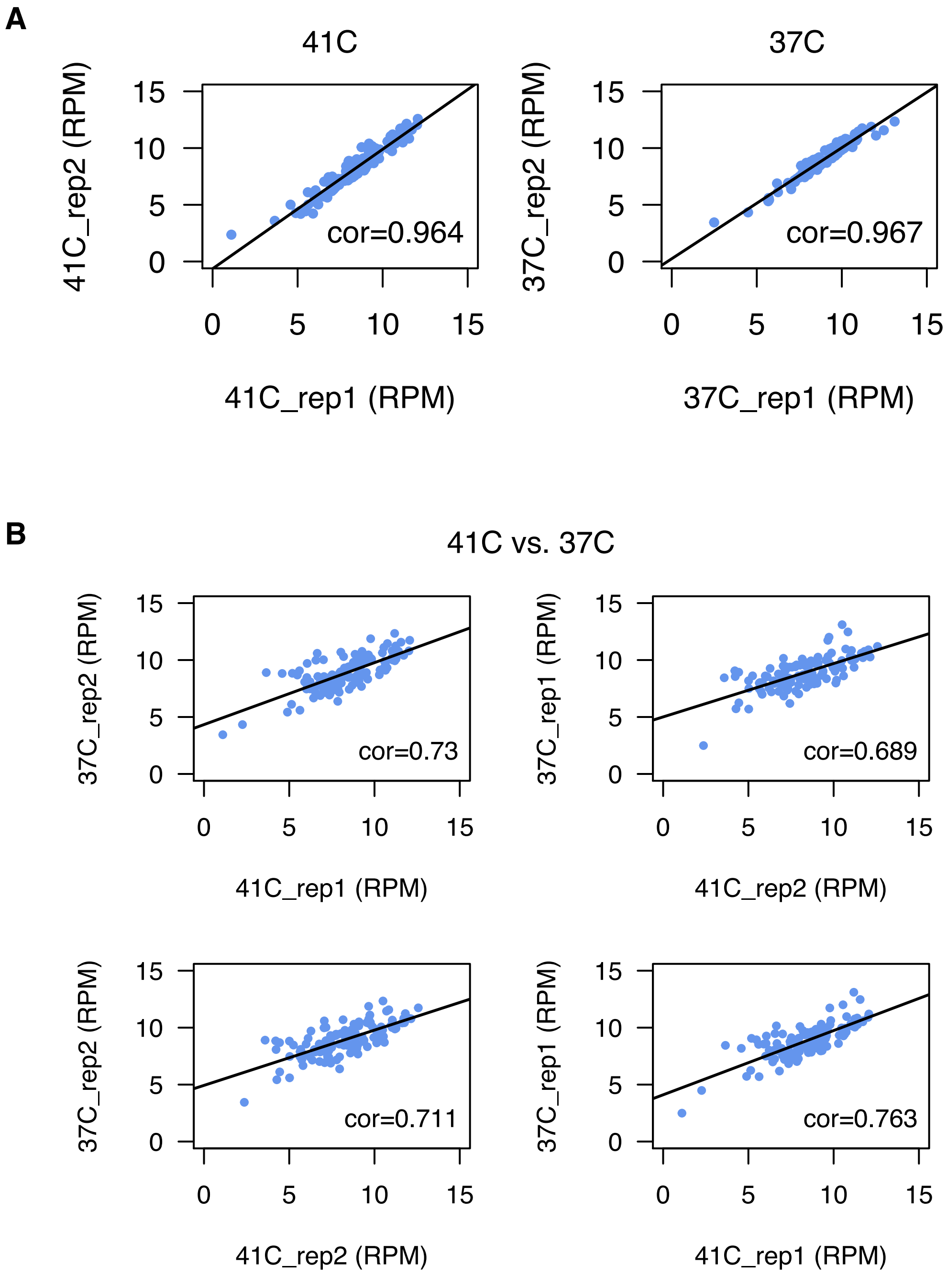

**Supplemental Figure S6**

**Randomness of *pB*-mutants' insertion-site distribution in the Pilot-Library and 1K-Library**

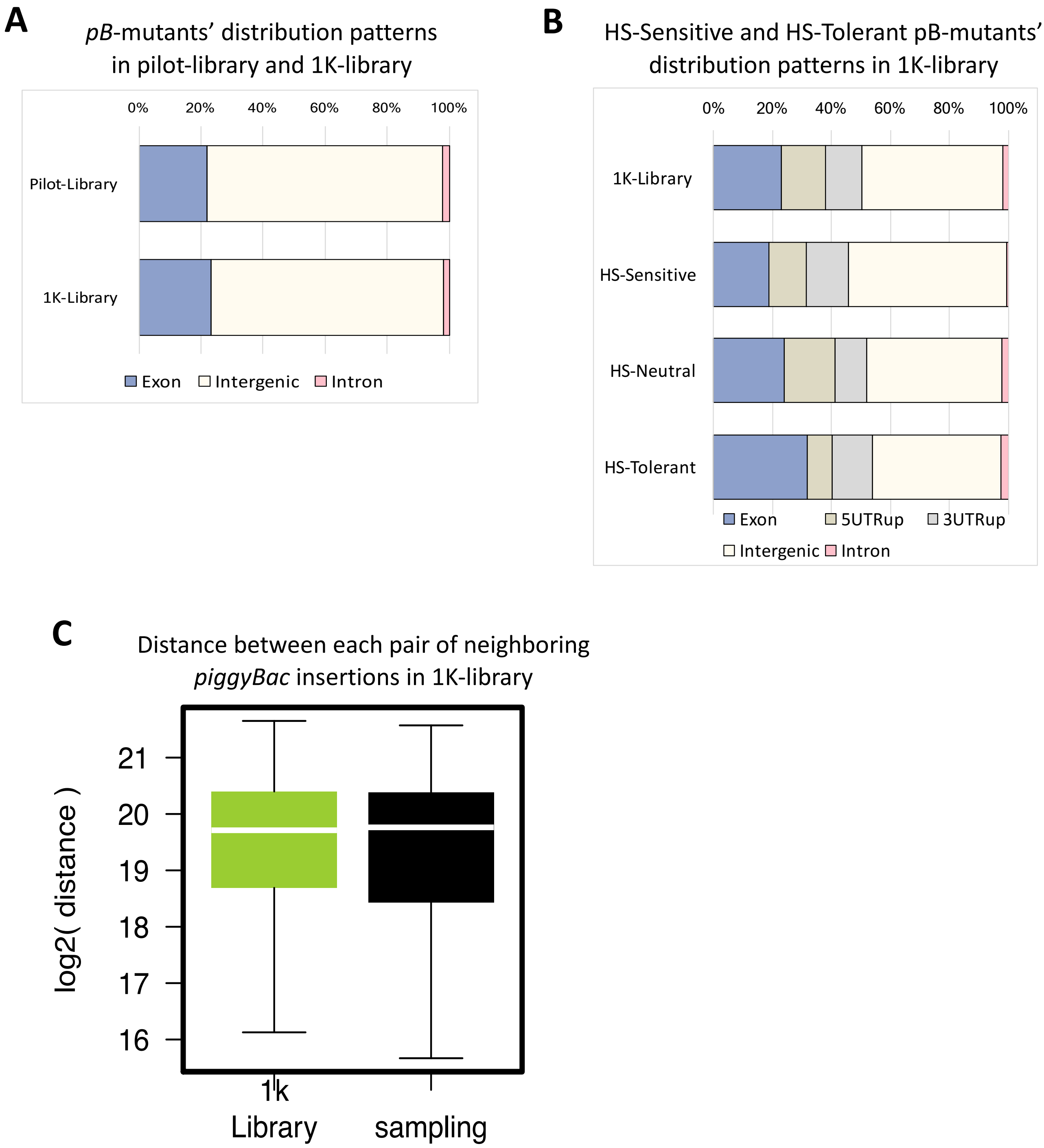

**Supplemental figure S7**

#### Reproducibility in the 1K-library HS-Screen

A

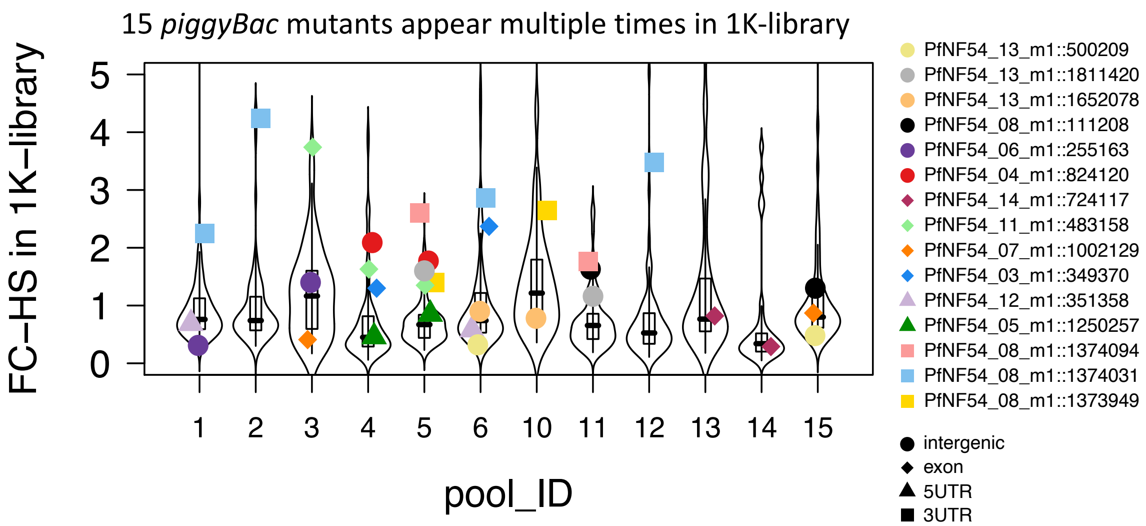


B

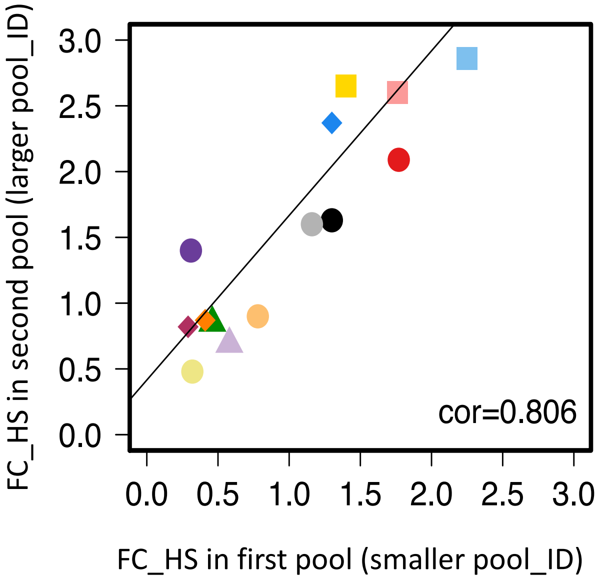


The Pearson correlation of 15 *piggyBac* mutants appear multiple times in 1K-library (cor = 0.806).

**Supplemental figure S8**

### Figure S8. Reproducibility within and between the pilot-library and 1K-library HS-Screens.

**A**

| PB Pool ID | PB Mutant ID | Insertion Site | Distance to gene | Type of Insertion | GeneID | Gene Function | Fold Change (41C/37C) |
|---|---|---|---|---|---|---|---|
| Pilot-library | PB4_DHC_11 | PfNF54_11_m1::879695 | 0 | exon | PF3D7_1122900 | dynein heavy chain, putative | 0.21 |
| 1K-library-Pool_14 | 14-016_DHC_12B | PfNF54_12_m1::135712 | 0 | exon | PF3D7_1202300 | dynein heavy chain, putative | 0.05 |
| 1K-library-Pool_4 | 4-054_DHC_10 | PfNF54_10_m1::980010 | 0 | exon | PF3D7_1023100 | dynein heavy chain, putative | 0.28 |
| 1K-library-Pool_14 | 14-015_DHC_12A | PfNF54_12_m1::121344 | 0 | exon | PF3D7_1202300 | dynein heavy chain, putative | 0.32 |
| Pilot-library | PB-54_FIKK9.3 | PfNF54_09_m1::99320 | 0 | exon | PF3D7_0902200 | FIKK family (FIKK9.3) | 0.52 |
| 1K-library-Pool_4 | 4-043_FIKK9.1 | PfNF54_09_m1::92390 | 1095 | intergenic | PF3D7_0902000 | FIKK family (FIKK9.1) | 0.23 |
| 1K-library-Pool_13 | 13-028_FIKK9.2 | PfNF54_09_m1::93353 | -477 | 3UTRup | PF3D7_0902100 | FIKK family (FIKK9.2) | 0.41 |
| 1K-library-Pool_12 | 12-040_ETRAMP | PfNF54_10_m1::82063 | 324 | 5UTRup | PF3D7_1001500 | early transcribed membrane protein 10.1 | 0.310 |
| 1K-library-Pool_1 | 1-055_ETRAMP | PfNF54_10_m1::82163 | 424 | 5UTRup | PF3D7_1001500 | early transcribed membrane protein 10.1 | 0.470 |
| 1K-library-Pool_4 | 4-048_HAD1 | PfNF54_10_m1::1339529 | -238 | 5UTRup | PF3D7_1033400 | haloacid dehalogenase-like hydrolase (HAD1) | 0.320 |
| 1K-library-Pool_12 | 12-037_HAD1 | PfNF54_10_m1::1340834 | 201 | 3UTRup | PF3D7_1033400 | haloacid dehalogenase-like hydrolase (HAD1) | 0.410 |

**B** Phenotypes of mutants in genes represented in both the pilot library and 1K-library (n = 16 genes) are highly correlated

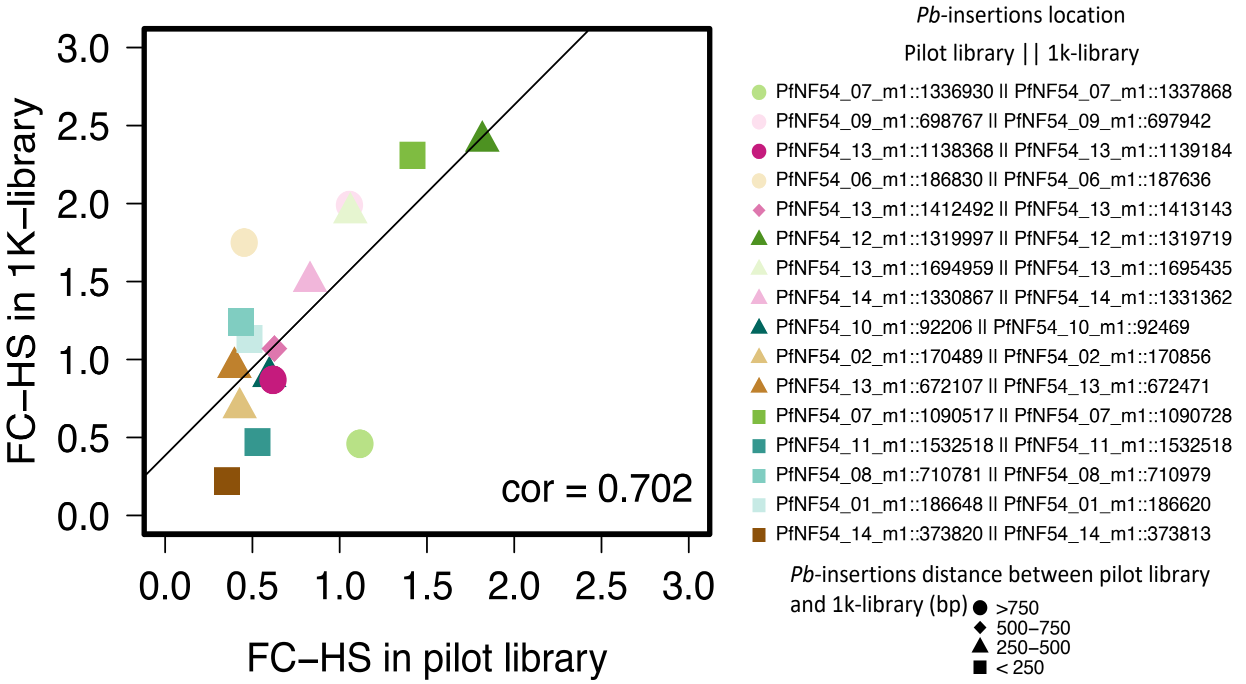

**Supplemental Figure S9**

#### Distributions of $PFS_{HS}$ for mutant heat-shock phenotype classifications in the 1K-Library screen

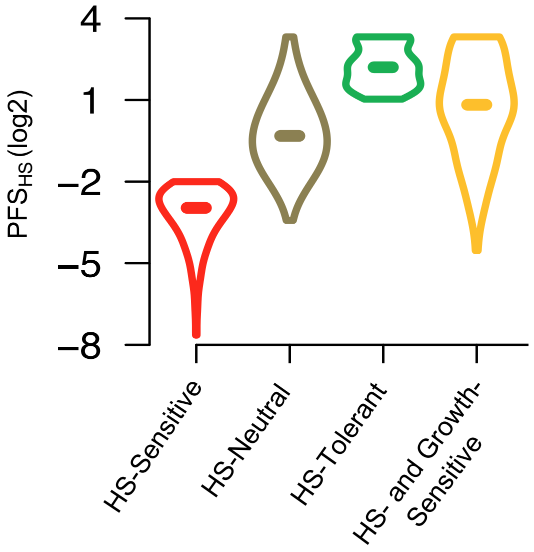

#### Supplemental Figure S10

##### QIseq data-correlations within and between Pilot-Library phenotypic screens

QIseq data Correlation of Bio-rep1 and Bio-rep2

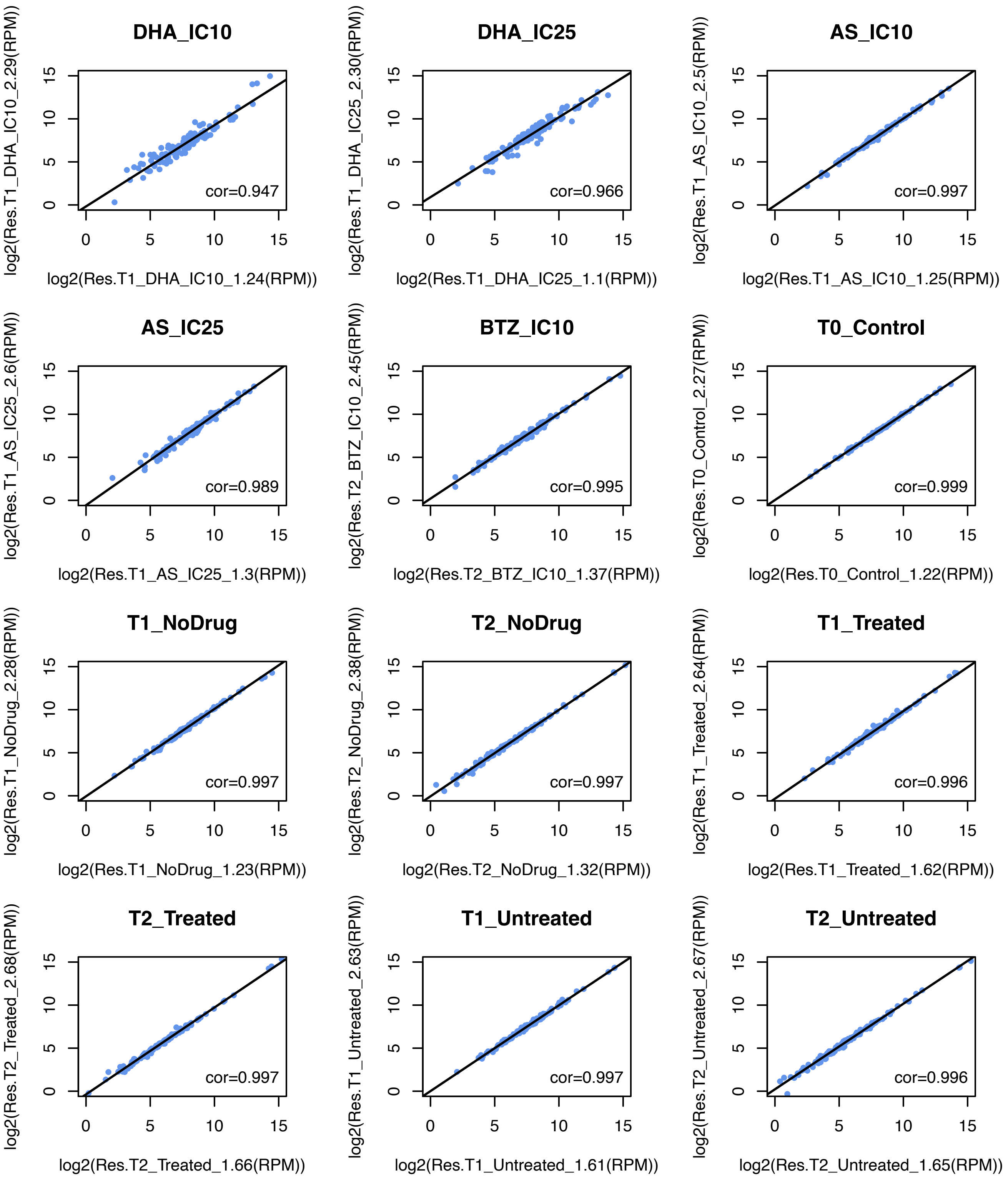

#### Supplemental Figure S11

### Validation of the HS-Sensitive phenotype mutations PB4 (*ΔDHC*) and PB31(*ΔLRR5*) during RNA-Seq Sample preparation

RNA sample collection for wildtype malaria-parasite NF54 vs. two HS-Sensitive *pB*-mutant clones in response to febrile temperatures

**A**

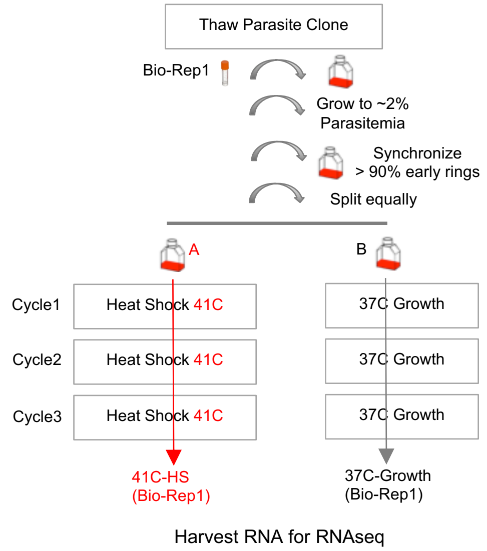


**B** Fold-change in HS vs. 37C-control

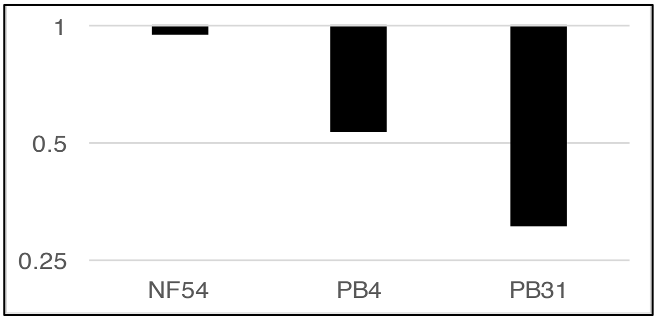

**Supplemental Figure S11**

**Complementary methods (pooled phenotypic screening, phenotypic transcriptional profiling of HS-Sensitive mutants vs. wildtype in response to heat stress) indicate genes driving the parasite heat-stress response**

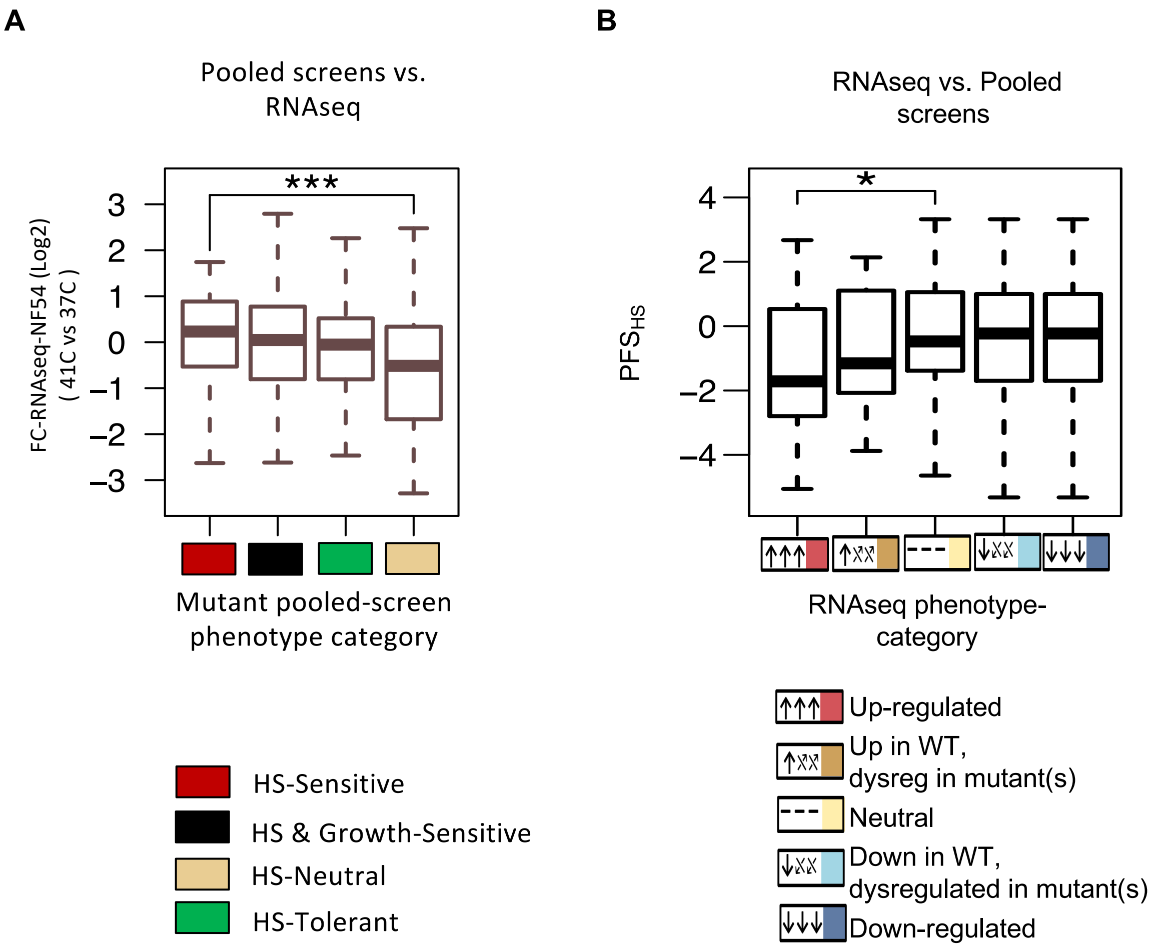
